# Improving Heart disease risk through quality-focused diet logging: pre-post study of a diet quality tracking app

**DOI:** 10.1101/2020.01.30.926634

**Authors:** Courtland VanDam, Bum Chul Kwon, Stephanie Chiuve, Hyung-Wook Choi, Paul Entler, Pang-Ning Tan, Jina Huh-Yoo

## Abstract

Diet-tracking mobile apps have been effective in behavior change. At the same time, quantity-focused diet tracking (e.g., calorie counting) can be time-consuming and tedious, leading to unsustained adoption. Diet Quality—focusing on high-quality dietary patterns rather than quantifying diet into calories—has shown effectiveness in improving heart disease risk. Healthy Heart Score (HHS) predicts 20-year cardiovascular risks based on quality-focused food category consumptions, rather than detailed serving sizes. No studies have examined how mobile health apps focusing on diet quality can bring promising results on health outcomes and ease of adoption. We designed a mobile app to support the HHS informed quality-focused dietary approach by enabling users to log simplified diet quality and view its real-time impact on future heart disease risks. Users were asked to log food categories that are the main predictors of HHS. We measured the app’s feasibility and efficacy on improving individuals’ clinical and behavioral factors affecting future heart disease risks and app use. We recruited 38 overweight or obese participants at high heart disease risk, who used the app for 5 weeks and measured weight, blood sugar, and blood pressure, HHS, and Diet Score (DS) measuring diet quality at baseline and the fifth week of the intervention. The majority used the application every week (84%) and significantly improved DS and HHS at the fifth week (p<0.05), although only 10 participants (31%) checked their risk scores more than once. Other outcomes did not show significant changes. Our study showed logging simplified diet quality significantly improved dietary behavior. The participants were not interested in seeing HHS, and the participants perceived logging diet categories irrelevant to improving HHS as important. We discuss the complexities of addressing health risks, quantity vs. quality-based health monitoring, and incorporating secondary behavior change goals that matter to users when designing mobile health.

## Introduction

An increasing number of mobile apps have explored ways to efficiently and effectively monitor and improve health behavior (1–3). Among these mobile health apps, diet monitoring is one of the most popular domains. Reasons attribute to diabetes and obesity leading as the top two fields producing revenue in the mobile health market (4) and their significance in impacting public health. A systematic review of mobile applications showed that mobile health apps increased adherence to diet monitoring and reduced efforts to maintain diet without using the apps (5). However, focusing on the quantification of diet can bring a number of challenges. Food journaling can be “too much effort,” “time-consuming,” or “tedious” (6,7). Detailed food journaling entry can be challenging because users often might not remember or know what and how much they have eaten (8). Users also feel the dietary information in the database is unreliable, calories burnt seemed random and “did not line up,” (9) and entering unhealthy food consumption in detail makes people feel guilty in general. These high barriers leading to limited engagement with diet tracking apps, researchers attempted lightweight approaches of diet tracking, and such attempt has shown to be successful by providing users a photo-based food tracking app and encouraging them to track only one food per day (10).

A 2018 study published at the Journal of American Medical Association showed the effectiveness of focusing on diet quality over quantity—to focus on restricting low-quality foods, such as processed foods, added sugar, or refined grains—rather than calorie counting (11). However, mobile apps on dietary monitoring focused on quantification of diet (e.g., calorie counting) and other health behaviors (e.g., steps). This quantification approach does not necessarily address the needs of broader groups of individuals. Numeracy and literacy in general can be a barrier. People show increased confusion around serving size (12), but for these apps to work appropriately, it would require accurate calculations of these very nuanced behavior choices. For instance, one might have eaten mixed salad, but the system needs to know how many grams of spinach versus carrots and which salad dressing in order to calculate accurate calories and nutritional content. Sophisticated, detailed, quantified tracking practices are not popular for all user groups (13). Tracking detailed health information is a user burden, affecting sustained tracking behavior (7).

At the center of effectiveness that mobile health brings includes seeing the effect of behavior change. Knowledge of risk level helps individuals to understand how urgent they need to change their behavior. Individuals at higher risk are more motivated to change if they know they are at high risk (5). A mobile app allowing users to observe how their risks are affected by their day-to-day choices relating to health and wellness (e.g., such as their choice of food that day) can greatly help with the prevention of chronic diseases. Awareness of heart disease risk has shown as one of the most critical methods and strategies to change behavior. Numerous mobile apps have been designed to directly or indirectly bring awareness about heart disease (14,15). However, these apps rarely show how lifestyle behavior change of risk factors—smoking, diet quality, or alcohol consumption—affects their outcomes to preventing heart disease (1,14–20). While understanding future risks increases motivation of individuals to change behavior, whether individuals will actually change behavior is a more complicated, sophisticated problem to solve than just “getting the message across” (21).

Our goal was to design and test a mobile app that would help users focus on improving diet quality with the help of getting real-time feedback on future heart disease risk as a result of their diet quality patterns. This way, we could increase individuals’ awareness on cardiovascular risks based on daily dietary choices. Users thus can focus on the behavior that is present and immediate, rather than an uncertain future (22,23). Users can log simplified categories that have high quality diet—e.g., vegetables, fruits, whole grains--to help users focus on the quality of food, rather than the detailed nutritional value, calories, and quantity of food.

Our study demonstrated that: (1) Monitoring simple diet quality can have significant effect on dietary behavior change; and (2) regardless of participants’ interest toward heart disease risk, the app reduced the risk.

## Materials and Methods

We designed the app based on Behavior Change Techniques (BCT) (24). We used focus groups to iteratively improve the paper prototypes and developed Android based app as a result. We then conducted a 5-week pre-post study with a follow up two weeks after the post study to evaluate the app’s efficacy of clinical and behavioral outcome changes as well as app usage patterns.

### Focus groups for app development

We conducted three focus groups in a sequence (n=13 total with 3~5 people for each group) to iteratively improve the initial digital paper prototype (Figure 1). The participants were at risk for heart disease recruited from a weight management clinic in the U.S. Midwest. During the focus groups, the participants were presented with images from the initial prototype to test usability and learnability of each screen (Figure 1). We revised the design iteratively based on the feedback. We then developed the mobile application on an Android platform.

**Figure 1.**
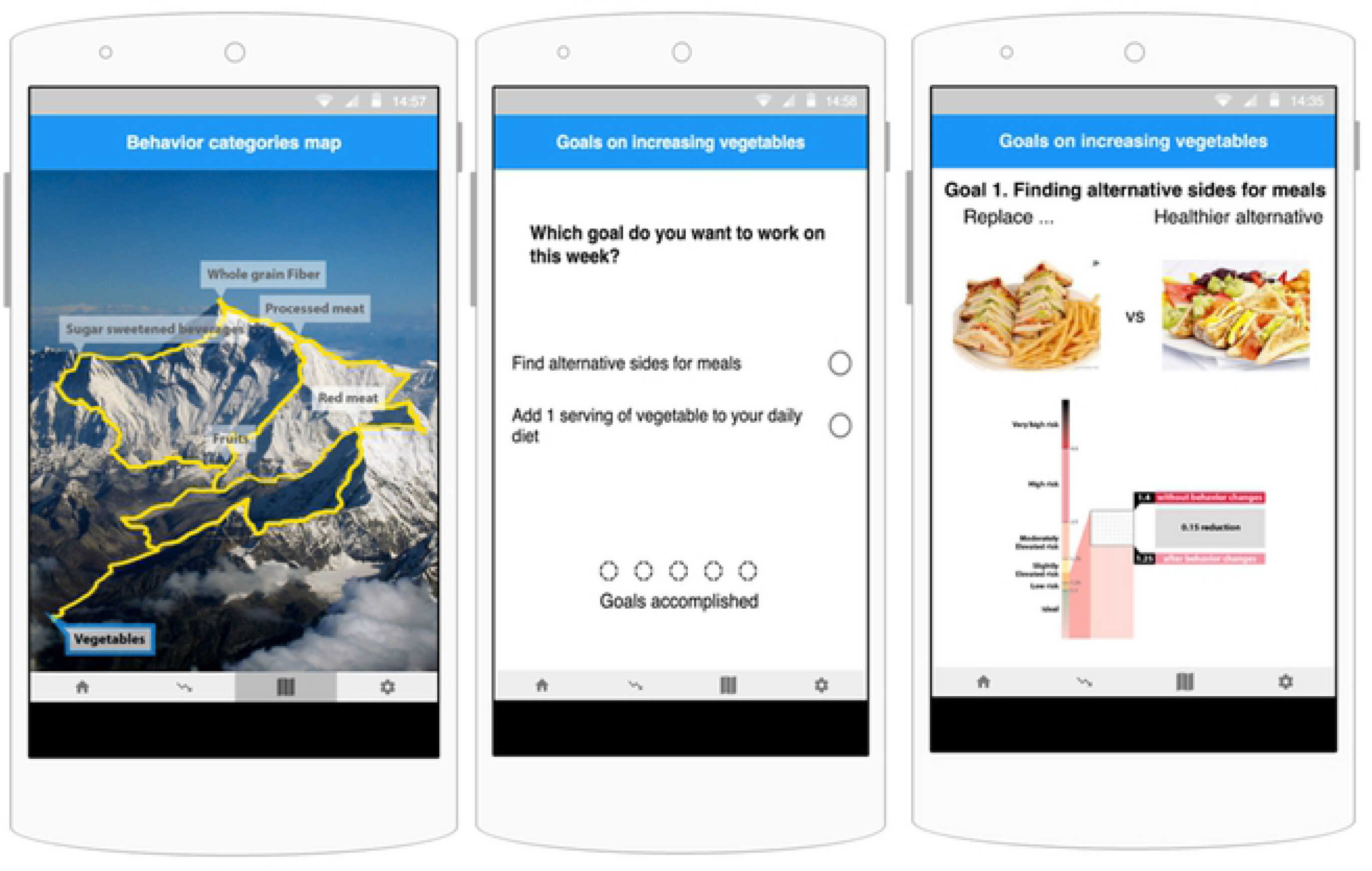
Screens from the prototype presented to the focus group. Users can select which goal to work on using the mountain climbing metaphor (left). As users accomplish the goals, they would unlock the next category of goals. Selecting a category on the Behavior Category Map would direct the user to the Goal Selection screen (middle). Right describes a screen in which users can choose the ‘sides’ and see how future cardiovascular risks might differ, if the user were to repeat the behavior for a week.

### Final app design

BCT suggests four core components to designing an intervention: Environmental contexts, Goals, Feedback and Monitoring, and Reinforcement. The app contains five screens: *Main Menu*, *Profile*, *Goals*, *Meal Calendar (food logging screen)*, and *Cardiovascular Risk (screen showing heart disease risk score)*. We designed the Profile page to incorporate environmental context, the Goals menu for users to personalize goals, Meal Calendar to log diet quality for feedback and monitoring, and Cardiovascular Risk screen to reinforce and reward positive diet change. A first-time user is directed to the Profile screen to provide their demographic information related to calculating their risk.

#### Diet quality and Healthy Heart Score (HHS)

The definition of high quality diet in this study was based on the Healthy Heart Score (HHS), a risk score system for heart disease risk developed at Harvard University (Figure 2) (5). Among a number of heart disease risk models (e.g., Framingham (25)), HHS is uniquely useful for middle aged adults who do not have elevated clinical factors, such as high blood pressure or cholesterol, but still may be at high risk for developing cardiovascular disease. The HHS model builds on lifestyle factors, such as smoking status, level of physical activity, alcohol intake, and a diet score based on consumption of fruits and vegetables, nuts, cereal fiber, sugar-sweetened beverages, and red and processed meats. HHS measures diet quality with the Diet Score (DS) factor (Figure 2). High DS indicates the individual is eating more *healthy foods*, including fruits, vegetables, nuts, and white meat. Consumption of *unhealthy foods*, including red meat, processed meat, and sugary drinks will lead to lower DS.

**Figure 2.**
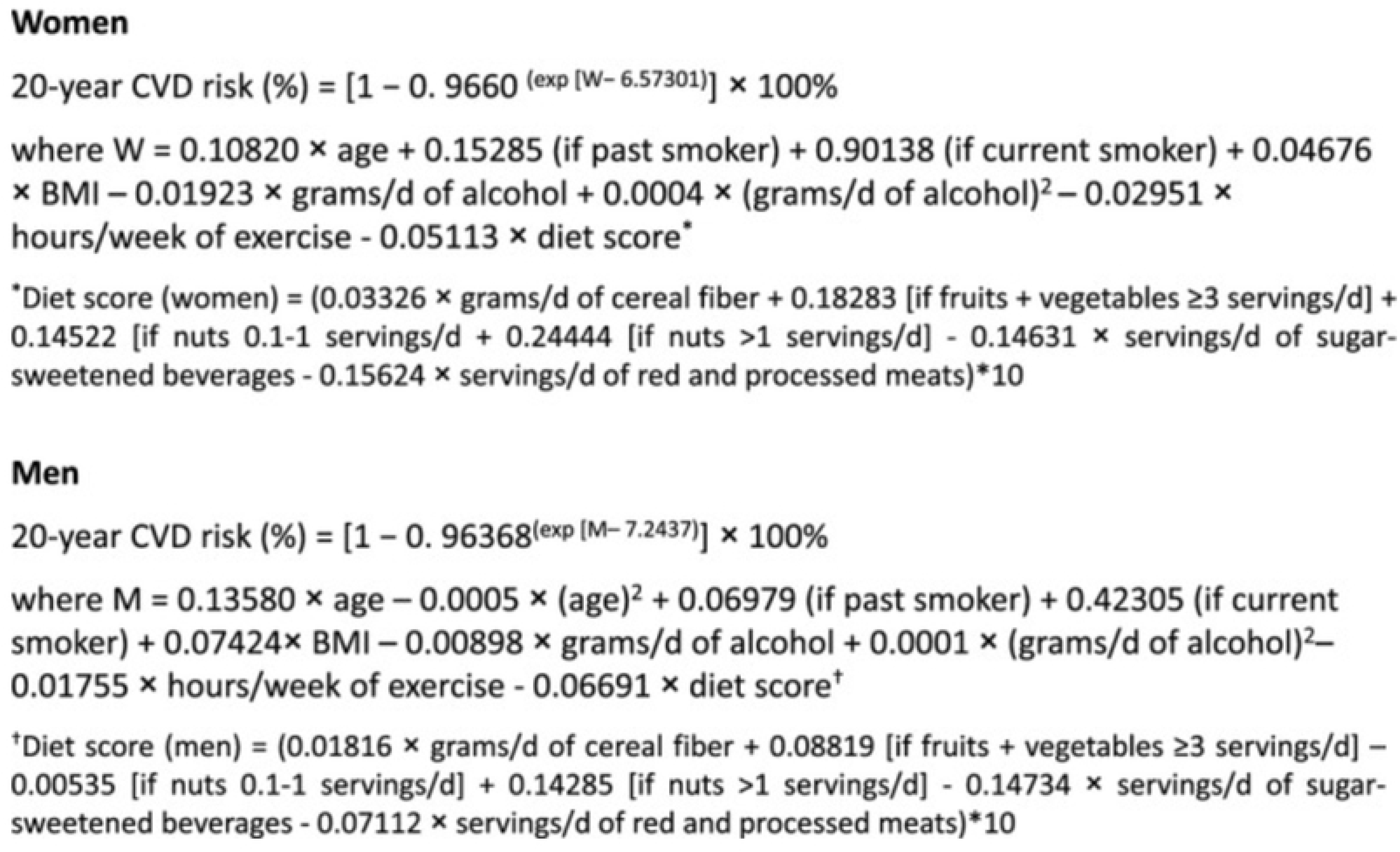
This figure describes the Healthy Heart Score (HHS) (5) and calculation of Diet Score (DS) for women and men.

In the diet monitoring screen (Figure 3a), users can enter up to four food categories for each of the meals they ate each day; breakfast, lunch, dinner, and snack. Following HHS, users could log overall quality of diet through the seven food category items noted by HHS: four *healthy categories*—fruits, vegetables, whole grains, and nuts and three *unhealthy categories*—red meat, processed meat, and sugary drinks. The app also allowed selecting the Other category to log foods not included in the provided categories. The Goals screen showed the default number of servings suggested for each food category. Users can either drag a food category icon, e.g. Fruit, to one of the meal slots, which counts as one serving of that category to that meal, or tap the calendar and work on the popup window to increase or decrease the number of servings and add the name of the food they consumed. Definition of a serving was not defined—any consumption counted as a serving, following the anti-quantification approach. In the Goal screen (Figure 3b), the default suggestions on the intake amount of unhealthy food categories was set to 0 servings. For fruits and vegetables, their combined total should be at least 3 servings per day, or equivalently, 21 servings per week.

**Figure 3.**
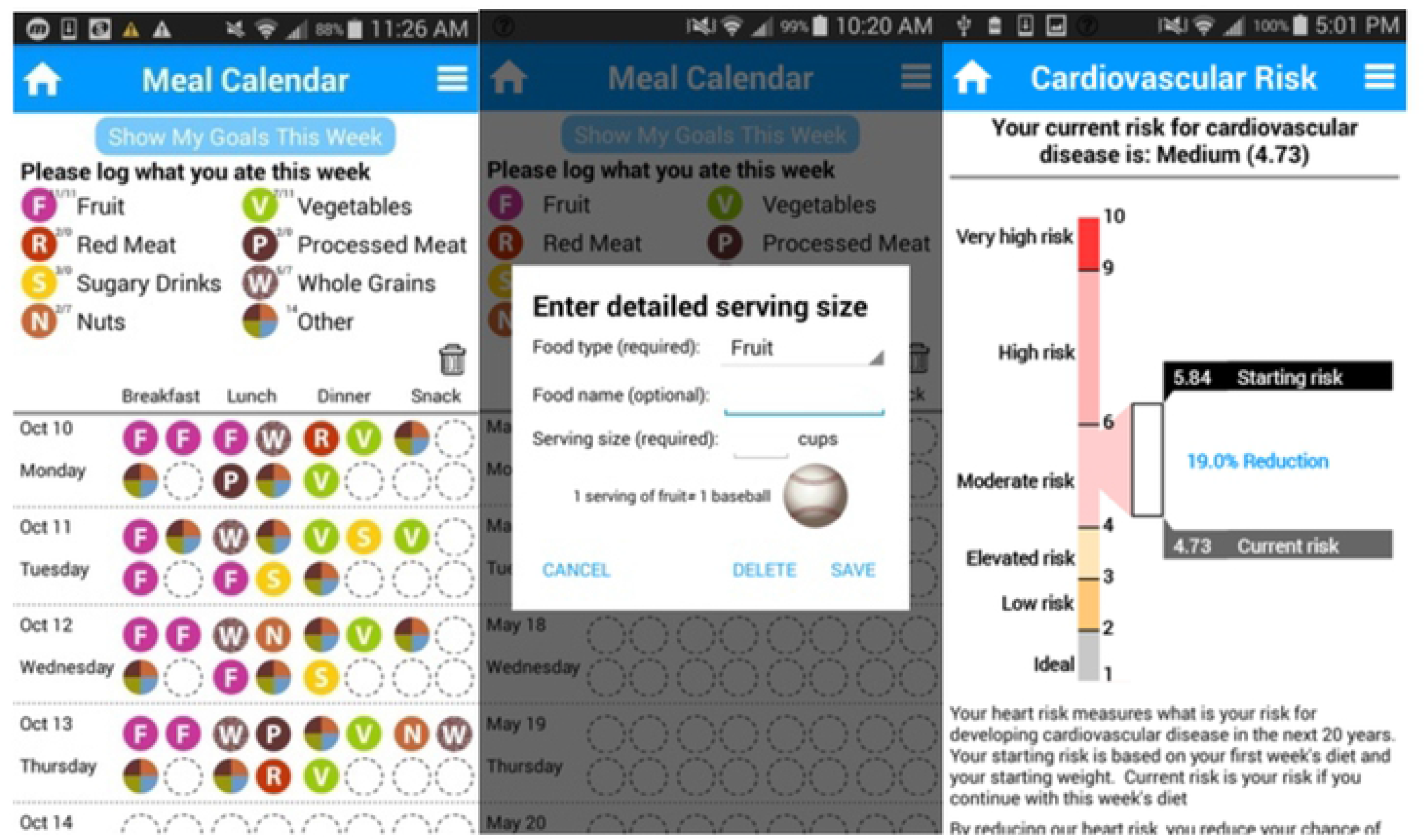
Figure 3a (left) shows the Meal Calendar screen, where users can enter simple quality-oriented diet categories. Figure 3b (middle) shows options to add more detail on the food, if user desires. Figure 3c (right) shows the screen that updates HHS risk score as user enters diet information.

#### Future cardiovascular risk

Cardiovascular Risk (Figure 3c) screen shows current HHS, the user’s real-time calculated cardiovascular risk score. The screen compares their risk when they started using the app to the current week. In this risk screen, we rescaled the HHS to a range from 1 to 10 from its original unit, 0-100%, following the suggestion provided by from the focus groups and in consultation with the expert who developed the HHS. The focus groups’ complaint was that the percentage was confusing—e.g., whether 50% meant 50% higher risk than others or half of the risk compared to others (or compared to current status). In the rescaled range of the score, the ideal risk score for a healthy individual is between 1 and 2, and if one has a risk score of 9 or above, the person is four times or more likely to develop heart disease than an individual with a healthy lifestyle.

#### Goals

At the beginning of each week, the app prompts users to set their goals and direct them to the Goals screen. Users can tap the goal icon of the food categories they want to actively work on. The users can deactivate a goal by tapping it again, and the goal will be displayed as grayed out. If one of the goals for unhealthy categories is active, users will be notified on the Meal Calendar if they exceed the maximum number of servings stated by the goal. The Goals screen includes a checkbox that shows the status of whether the user has met the goal.

### Pre-post study: Recruitment and procedures

The participants were recruited from a weight management clinic at a major hospital in the U.S. Midwest. 38 participants started the study between June and September 2016 for a 5-week intervention (denoted as Week 0-4) with a follow up meeting two weeks after the end of Week 4. The participants were asked to use the app at least six days a week for the five weeks of the study. The participants had the option to continue using the app until the follow up meeting. Initially, the participants were asked to log their diet to establish a baseline. Starting at the beginning of Week 1, the app started prompting the participants to set goals for each week based on HHS recommendations--either by keeping the default suggestion (ideal diet) or changing it to personalized goals.

At baseline and at the end of Week 4, the participants visited the clinic for a clinician to measure their weight and fasting blood sugar. At the end of Week 4, we reminded the participants they were no longer required to use the app. In addition, an exit interview was held at the follow up visit to discuss their experiences with the app. Figure 4 shows the study procedure.

**Figure 4.**
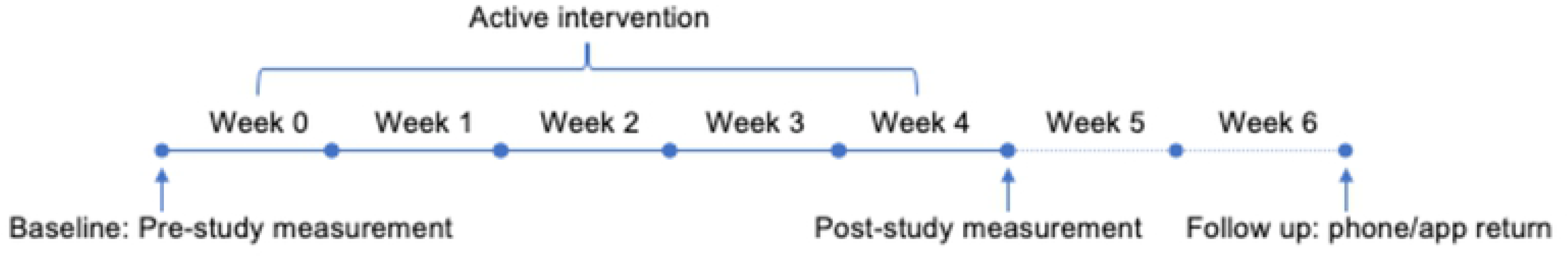
The figure shows the timeline of the pre-study and post-study measurements and follow up and the notation of the Weeks.

All participants received cash compensations of up to U.S. $50. The participants received partial or full compensation depending on how much they completed the following: three online surveys; measure health outcomes twice; and use the app at least 6 days a week during Week 0 to 4. The app was provided to the participants in two ways. If the participants had an Android phone, we installed the app onto their phones. Otherwise, we provided them with a Samsung Galaxy S3 phone, with the app installed, for the duration of the study. These participants were required to return their phones at the follow up.

### Research questions and analyses

We wanted to answer three research questions regarding feasibility to logging diet quality, motivating behavior change through feedback on future heart disease risks, and the app’s efficacy of behavior change.

### RQ1. How feasible was logging diet quality?

We recorded and analyzed the time, frequency, and screen of the participants’ tapping events on the app. We analyzed how often participants went to each screen and which food categories were logged over time. We also analyzed user logs about food names to understand diet logging behavior.

### RQ2. How feasible was communicating risk to motivate behavior change?

We analyzed participants’ usage of the Risk screen. We then associated the usage with participants with their HHS.

### RQ3. How effective was the application in changing health outcomes?

We conducted a paired-sample t-test to compare the outcome changes in diet quality, HHS, and in-clinic measurements (weight and fasting blood glucose) between pre- (At the beginning of Week 0) and post-test (At the end of Week 4) measurements.

## Results

In this section, we first report demographical information of the participants and the recruitment outcome. We then report results on diet quality, risk score checking, health outcomes and diet score, and association between app use and diet score.

### Participants

32 out of the 38 recruited participants completed at follow up. Twenty-two participants used the provided study phones and the rest used their own phones. Three participants who were Android phone users stopped using the app and stopped responding to the researchers. Another participant withdrew because the app was too cumbersome. Two other participants withdrew because they decided the study no longer applied to them. The remaining 32 participants (Female=26; Age: M=57.48, SD=11.85) who completed the study have a wide range of age, weight, smoking status, and experience of using a smartphone. One participant was a smoker, 12 were former smokers, and 19 were non-smokers. 17 participants were diagnosed with diabetes. Four participants were overweight (BMI between 25 and 29.9) and the remaining 28 participants were obese (BMI >= 30) (26).

### RQ 1: How feasible was logging diet quality?

#### App use

During the active intervention when the participants were required to use the app (Baseline~Week 4), the participants tapped on the app 27 times on average (SD=25.6) per week. After Week 4 and until follow up, the participants tapped on the app 11 times on average (SD=18.3) per week. Figure 5 shows each participant’s overall app use over the weeks.

**Figure 5.**
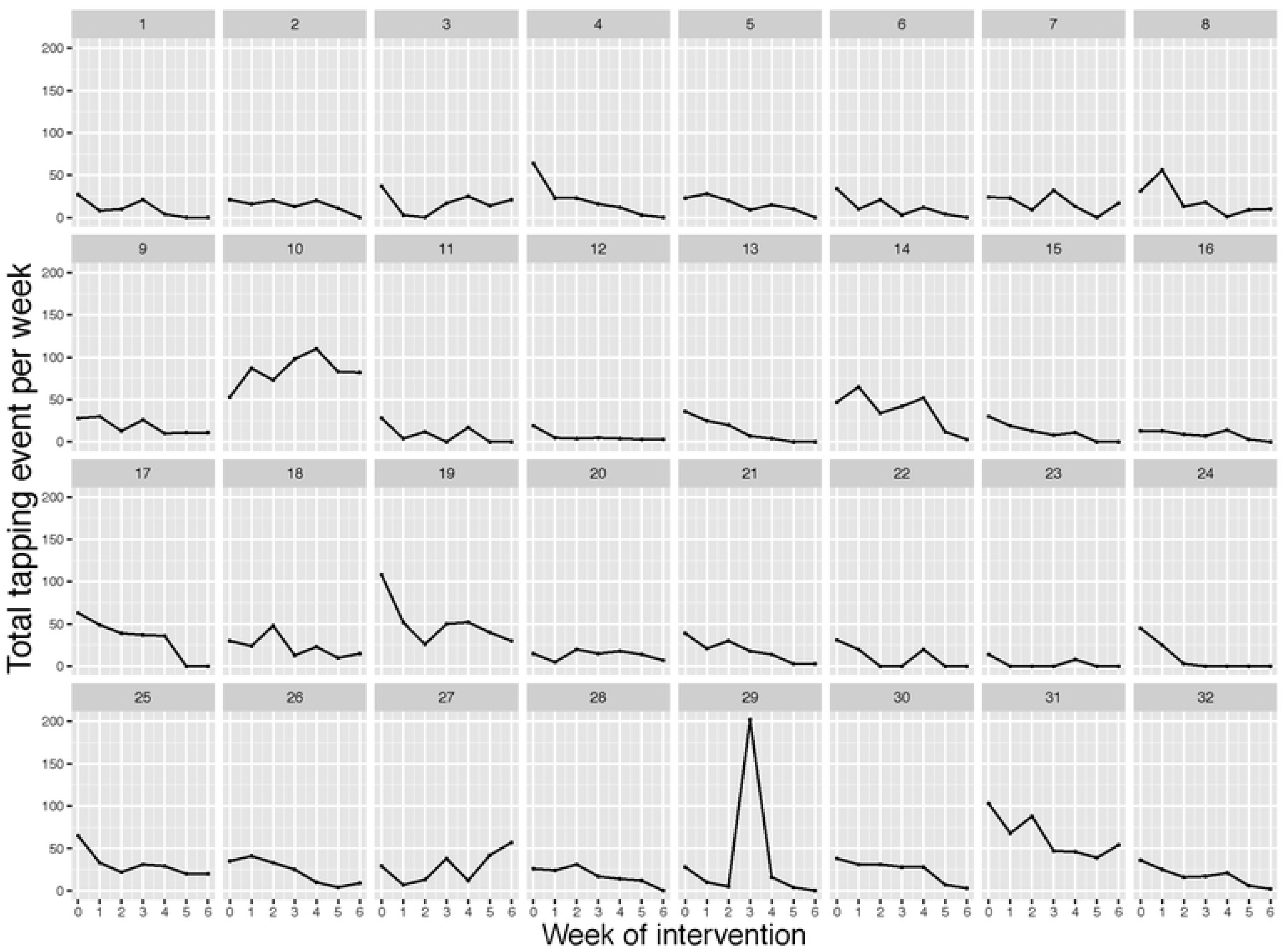
Each small graph shows each participant’s total number of tapping events over the 7 total weeks including two-week follow up (between the baseline and follow up). The x-axis shows the week of intervention (0 indicating the frequency accumulated between the baseline and at the end of Week 0). The y-axis shows the total number of tapping for each week. 27 participants visited the screen nearly every week for the intervention duration (Week 0-4).

#### Diet logging

28 out of 32 participants logged food every week between the baseline and at end of Week 4, at least once a week. About a half of the participants logged food nearly every day. As seen in Figure 6, during the active weeks, the 32 participants altogether logged “Other” the most (6066 instances, 40%), followed by “Vegetables” (2857 instances, 19%), “Fruit” (2398 instances, 16%), “Whole Grains” (1614 instances, 11%), “Nuts” (698 instances, 5%), “Red Meat” (627 instances, 4%), “Processed Meat” (614 instances, 4%) and “Sugary Drinks” (151 instances, 1%). Figure 6 shows when these food categories were logged to the meal slots during the course of the day—breakfast, lunch, dinner, or snack. Fruits and whole grains were logged proportionally larger during breakfast meals than other meals, and vegetables were logged proportionally larger during lunch and dinner meal times. “Other” categories were logged equally over all meal slots.

**Figure 6.**
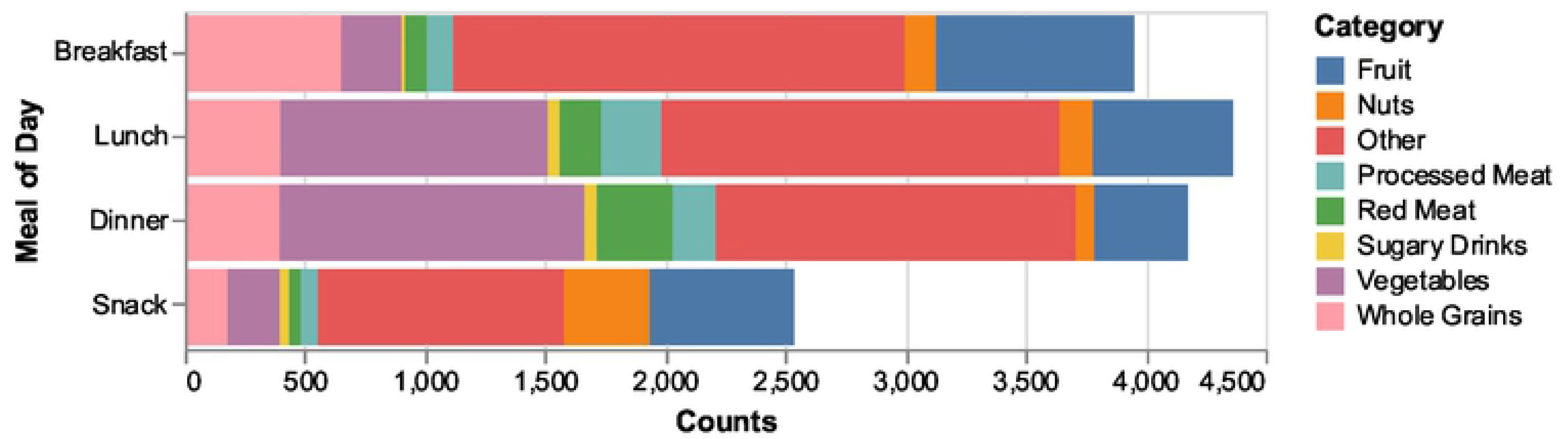
This figure shows all 32 participants’ logging per food category and which meal of day the logging occurred during the active intervention weeks (Between baseline and end of Week 4).

Participants entered qualitative description of the food in the “food name” field for 38% (5,730 instances) of all diet logging instances. 49% of these instances (2,800) were entered when logging to the Other category, 18% for the Vegetables, 15% for Fruit, 8% for Whole grains, 4% for Nuts, 3% for Red meat, 3% for Processed meat, 0.7% for Sugary drinks. For non-other categories, participants entered example description of the food category they entered. For instance, Fruit category included descriptions such as “strawberries” or “grilled fruit salad.” Vegetable category included “arugula” or “grilled squash and zucchini with lemon and olive oil.”

When participants entered “Other,” 98.9% of them included detailed descriptions on the food. The qualitative analysis of these descriptions together with the exit interviews revealed several reasons for why the food was logged an “Other.” First, the given food categories did not capture all the food categories they attempted to log, such as their current dietary goals (e.g., to reduce dairy). The participants were given the instruction to only log what is related to heart disease risks, but they still captured other categories not affecting healthy heart risk, including dairy, dessert, or other protein foods (e.g., 338 protein instances such as eggs, tofu, and beans, 12.2%; 584 dairy instances such as milk, cheese, and greek yogurt, 21.1%). The participants also captured foods in the “Other” category when the food was a mix of various food categories that might have been difficult to be captured in one or two food categories (e.g., California roll, sandwich). 19 instances showed red meat food, such as pork, and vegetables being logged as the “Other,” showing how the users might have been confused on what food categories these foods belonged to. Even though pork was red meat, the fact that it was logged as the “Other” matched with the exit interview content that the participants considered pork a white meat.

### RQ2. How feasible was communicating risk to motivate behavior change?

#### Risk screen

As Figure 7 shows, at the baseline, most participants checked their Risk scores (n=29). Starting the week after, however, most did not come back to the Risk screen to view their changes in their HHS. Thirteen people checked the Risk screen in Week 1, 11 in Week 2, 6 in Week 3, 10 in Week 4, and 6 in Week 5 until follow up.

**Figure 7.**
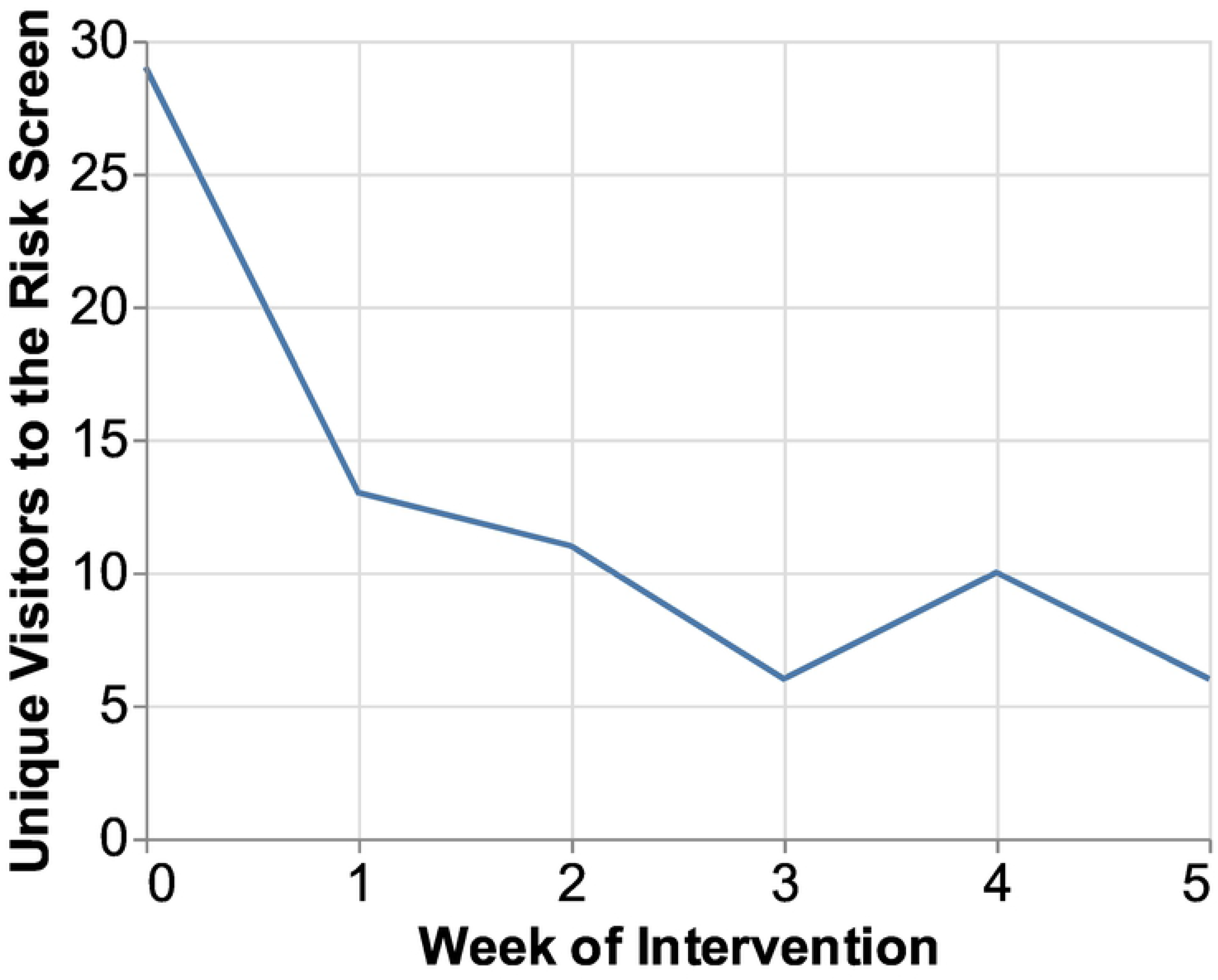
The figure shows the participants’ use of the Risk screen (loading frequency) over the weeks. 29 participants out of 32 checked their risks the first week, and then only a few checked again at Week 4 (n=10). Most participants did not return to the Risk screen to recheck it after the baseline.

### RQ3. How effective was the application in changing health outcomes?

#### Diet Score

All but two participants logged their diet during the active intervention. Among the n=30 participants who logged their diet during the active intervention, the Diet Score showed significant difference between baseline (M=1.31, SD=1.14) and post-test during Week 4 (M=2.36, SD=2.48); t(29)=-2.85, p=0.008. (See Figure 8).

**Figure 8.**
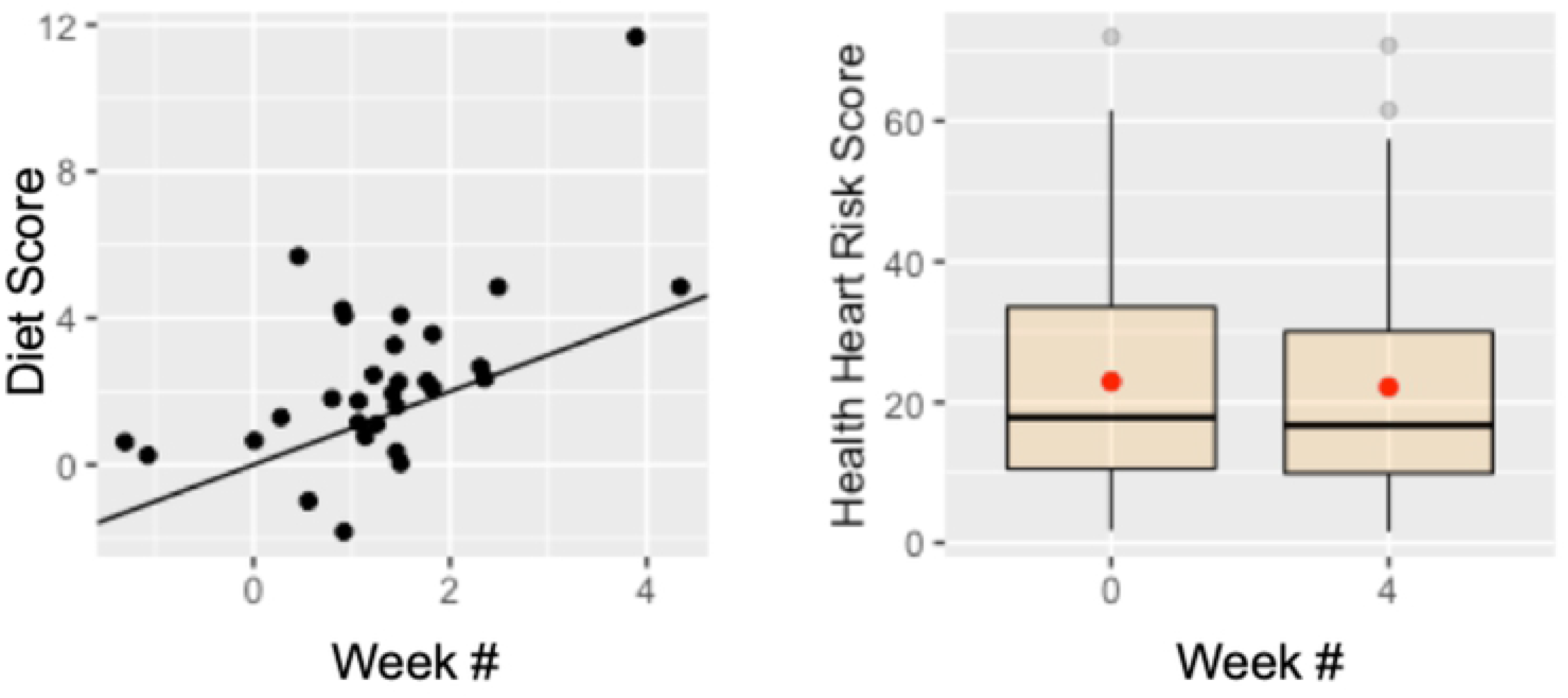
The figure shows the Diet Score (left) and Healthy Heart Score (right) changes between pre- and post-study measurements of the participants.

#### HHS

Healthy Heart Score also showed significant difference between baseline (M=22.94, SD=18.86) and post-test at the end of Week 4 (M=22.15, SD=18.58) measurements; t(29)=2.41, p=0.02.

There was no statistical association between food logging frequency and three measures: Diet Score, Risk, and BMI.

#### In-clinic measurements

Weight did not show significant difference between pre-test (M=241.7 lbs, SD=61.17) and post-test (M=242.6 lbs, SD=61.9) measurements; t(29)=-1.043, p=0.31. Blood sugar also did not show significant difference between pre-test (M=130.2, SD=76.62) and post-test (M=123.3, SD=48.8) measurements; t(28)=-0.95, p=0.35.

## Discussion

The study showed feasibility to logging diet quality (RQ1) but not communicating risk (RQ2). However, the application was effective in changing health outcomes (RQ3), showing logging simplified diet quality significantly improved dietary scores and future cardiovascular risk scores. The following shows key takeaways:

- The study showed no association between frequency of logging and improved dietary scores, showing the importance of separating frequency of use in measuring health outcomes.
- The participants were not interested in monitoring the risk scores, but they still significantly decreased their risk scores by focusing on the target behavior. This finding gives implications to health risk communication in mobile health app design.
- The study showed users mostly logged irrelevant dietary behaviors to the target behavior. This finding shows the need to balance reducing monitoring items for efficiency versus what matters to users to support user experience.

### Opportunities and challenges of quality focused diet monitoring

Previous literature shows logging diet is highly associated with improved diet (27). At the same time, studies showed that not all users can benefit from sophisticated diet logging applications. Users often find diet logging a tedious, cumbersome activity, which leads to abandonment (6). Also, people do not always accurately estimate food proportions and nutritional contents (12). Automated techniques including calorie calculations and artificial intelligence-based food detection can reduce such user burden in detailed diet logging (28–30). However, these methods are still limited and error prone, which lead to increased user frustration and abandonment.

To address this gap, we implemented the Healthy Heart Score (5) into a mobile app, which simplified the diet monitoring process to focusing on improving diet quality over quantity. This approach incorporates a lenient approach toward food proportion and nutritional details in calculating the risk. By allowing users to focus on logging simplified diet quality that does not require logging detailed nutritional, caloric breakdown of each meal and focusing on whether a gross food group was consumed (fruits, vegetables), we showed users steadily used the app even after the required weeks they were not incentivized to use it. One participant asked if they can continue using the app even after the study had completed.

At the same time, the study showed no association between frequency of use and diet score increase. This finding shows the need to separate quantitative measure of usage from health outcomes. This implication aligns with the discussions around whether sustained use of an mHealth app is a positive one or not—discontinuing to use an app might mean the user no longer needs the app because the user has achieved the health goal or that the user has become more independent (9).

One challenge we discovered in logging diet quality was that even at the gross level of food categories, some participants found confusions around categorizing food to the right categories (e.g., confused pork as white meat, avocado as not being vegetable).

### Implications for health risk communication in mHealth design

Our initial goal of this app was to increase individuals’ awareness on cardiovascular risks based on daily dietary choices. We expected that users would check on their risk scores as they changed their dietary patterns to understand how their risks were impacted by their dietary choices, thus making behavioral changes. However, while logging the diet quality was positively accepted by the participants, the participants rarely visited the risk screen throughout the weeks. The participants mainly visited the risk screen at the very beginning to check their initial risk score, and a few came back for a second check after a week, and most did not come back. The follow up interview revealed that the participants noted their score did not seem to visibly change, so they did not think to check more often. At the same time, the HHS results showed that the participants still significantly improved their HHS at Week 4. A solution would be to improve on visualizing the risk scores so that their improvement is more visible and concrete. One idea is to augment a forecasting trajectory to the risk score. The predicted line could be designed to adjust more sensitively to users’ recent efforts to provide further motivation.

Communicating future risks is known to alert and motivate people to change behavior (31–33). At the same time, risk communication largely suffers from people making the actual behavior change because the risk is too distant in the future, giving lack of sense for relevance (34,35). This study showed the participants were initially motivated by their risk score, but the behavior change was not related to their checking of the risk score over time. Though users did not check their risk scores, they overall decreased the risk scores in the end. This finding gives implications to the role of health risk communication in consumer facing mobile health apps, in which continuous monitoring is the strength. The risk scores can serve as initial motivation to set up goals, but users would focus on monitoring and improving the target risk behavior (in this case diet quality), and the improvement with the risk can be a positive side effect.

### Implications of “Other” in monitoring apps

The findings on the largest logging activity of “Other” food categories provided implications for balancing between simplification and accommodation of users’ “Other” needs. The HHS discourages or encourages certain food categories to be consumed. This instruction—to focus on improving consumption of certain food categories—was reassured to the participants during the instruction. In the app design, we also specifically only allowed users to log the relevant food categories to improving the HHS score. However, the majority of the diet logs were under “Other”, where it included irrelevant food categories, such as dairy. According to follow up interview, this came from having a concurrent diet goal of their own. When designing a monitoring app to improve a health behavior, one needs to consider the gap between the chosen clinical approach and individuals’ concurrent goals and considerations. While simplifying the design to only monitor necessary information can improve efficiency and reduce user burden, this design approach might lose incorporating users’ concurrent needs and focus. One should not consider what matters to users as “Other” because it is considered irrelevant to a target goal.

## Limitations

This study did not have a control group, and the duration was only five weeks—not enough to show true behavior change. The data did not include collecting information on whether the participants did not continue to check risk scores because of the lack of usability (e.g., legibility of the visualization) or their disinterest on risks.

## Conclusions

Our study showed feasibility and efficacy of a simplified diet quality monitoring in a mobile health application. Future work should further test the app’s efficacy with a larger, focused population who are disinterested in using existing quantity-focused monitoring applications. Despite some known limitations on research design and duration, the findings provided significant contributions to understanding the implications on the opportunities and challenges in designing a simplified, diet quality focused monitoring app and how health risk communication can be effectively integrated into an mHealth design. The study also sheds light on finding the balance between affording users to focus on simplified target behavior, reducing user burden versus further incorporating what matters to users in designing a health monitoring app.

## Acknowledgements

None

## Supporting Information

